# Genomic correlates of metastatic competence and progression in human melanoma

**DOI:** 10.64898/2026.09.23.753695

**Authors:** Eftychia Chatziioannou, Karan Luthria, Parin Shah, Sorin Armeanu-Ebinger, Jakob Admard, Christopher Schroeder, Stephan Forchhammer, Frederik Stihler, Xingpei Zhang, Synaida Maiche, Zeynep Cakmak, Anette Barbara Mankel, Josua Stadelmaier, Teresa Amaral, Ulrike Leiter, Evelyn Maczey, Sven Mattern, Irina Bonzheim, Falko Fend, Claus Garbe, Yvonne Heneka, Marie Krauss, Oltin Tiberiu Pop, Laura Schreieder, Sebastian Haferkamp, Alexander J. Stratigos, Christian M. Schürch, Stephan Ossowski, Peter Martus, Sven Nahnsen, Tobias Sinnberg, Lukas Flatz, Olaf Riess, Benjamin Izar, Martin Röcken

## Abstract

Genomic events and their timing that grant a primary tumour the competence to disseminate remain poorly defined. We performed sequencing of 247 stage I/II primary cutaneous melanomas (CMs) and 60 matched metastases without intervening therapy from a prospectively followed registry cohort with a median followup of 92 months, integrating copy-number, mutational, protein and spatial-transcriptomic analyses. Relapse was not distinguished by oncogenic point mutations, which were largely shared between primaries and metastases, but by somatic copy-number alterations (SCNAs) and global chromosomal instability. We defined OncoCycle, a six-gene copy-number signature (amplification of *CDK4*, *MCL1* and *CD276*; biallelic loss of *CDKN2A*, *CDKN2B* and *TP53BP1*) that predicted relapse independently of established clinicopathological features in melanoma, and a pan-cancer analysis. In matched pairs, metastatic progression was driven by continued copy-number evolution and reduction in intra-tumoural heterogeneity, rather than by acquired point mutations, and OncoCycle alterations from primary tumours were preserved in metastasis seeding clones. Clonal reconstruction revealed both monoclonal and polyclonal metastasis seeding, and spatial transcriptomics resolved copy-number-defined metastatic subclones occupying and programming distinct immune and stromal niches. Thus, metastatic competence was primed early by focal SCNAs on a background of chromosomal instability, elaborated by continued copy-number evolution during dissemination and spatio-temporal interactions with the tumour-microenvironment.

## Introduction

While stepwise genetic emergence and evolution are well characterized in carcinogenesis of primary tumours, mechanisms of metastasis in patients remain remarkably poorly understood^1–3^. In the last century, different models of metastasis have been proposed: linear progression, where a multi-step acquisition of genetic factors selects for a clone with metastatic competence; parallel progression, wherein metastasiscompetent cells disseminate early and co-evolve with the primary tumour before the latter may become clinically apparent; and, metastatic randomness, which proposes an estimation of whether primary tumour subclones are selected for dissemination at random as compared to a quantifiable fitness acquisition that drives metastasis^4,5^. Two questions remain unresolved: whether metastatic competence is an intrinsic property established early in a tumour’s life or a capacity that evolves stepwise, and, if a heritable trait exists, what its molecular identity is. Somatic copy-number alterations (SCNA) have been proposed as a candidate marker, but direct evidence from matched, treatment-naïve human primary tumours and metastases is extremely scarce^1–3^.

Early-stage melanoma represents an instructive disease context to examine principles of metastasis^6^. The genomic evolution of melanocytes into cutaneous melanoma (CM) is well characterised^7,8^, whereas the genomic and microenvironmental factors that drive progression from primary into metastatic disease remain poorly understood^2,3,9,10^. Genetic, experimental and histological analyses of melanomas, and other cancers show that malignant cells can disseminate early and migrate to distant organs^11–18^; yet most CM never cause metastases, or do so only after years^19,20^. Panel and whole-genome sequencing of diverse and well-controlled patient populations, including melanomas, identified chromosomal instability as a recurrent feature of metastatic cancer, while driver mutations rarely diverge between primary tumours and metastases, consistent with other tumour types^13,21–28^.

A major obstacle in examining principles of metastasis in CM and other cancers is the collection of matched biopsies of primary tumours and subsequent metastasis collected *without* intervening therapy that may alter salient biological insights^1^. This is due to several factors, including in part long metastatic latency, fragmentation of care and lack of molecular profiling of the early-stage primary tumours, which is frequently clinically (and therapeutically) inconsequential. Here, we have overcome these challenges through a long-term, prospective collection of primary tumours, long follow-up without intervening therapies, and biopsies of matched relapsed metastatic lesions. Comprehensive copy-number, mutational, protein and spatial-transcriptomic analyses reveal different mechanisms of metastasis and clone-specific interactions with the tumour-microenvironment. We find that SCNAs, concentrated in a small group of genes governing cell-cycle progression, chromosomal stability and immune checkpoint B7-H3 (encoded by *CD276*), rather than point mutations, distinguish primary tumours destined to relapse, and that metastases arise from clones already bearing these alterations and subsequently acquire additional SCNAs in the same and in immune-related pathways. As loss of *CDKN2A* or activating *CDK4*-mutations prime localized melanocytic tumours for metastatic progression into melanomas also in mice^29–31^, metastatic competence in melanoma is primed early by acquisition of distinct SCNAs and elaborated through continued copy-number evolution during dissemination.

## Results

### Clinical and genomic features of study population

To identify genomic alterations that might predispose primary CM to metastatic progression, we performed targeted panel sequencing of cancer-associated genomic regions at a mean coverage of 385× for tumour samples and 290× for matched normal tissue (Table S1). We analysed 37 stage I (15%) and 210 stage II (85%) primary CMs, together with 60 corresponding metastases (Fig. 1A), from 247 patients enrolled prospectively in the German melanoma registry and diagnosed between 2000 and 2018, prior to the approval of effective adjuvant therapy with either immune checkpoint blockade or MAPK directed targeted therapies^32,33^. For each tumour we assessed histology, nucleotide variants and SCNAs.

**Figure 1.**
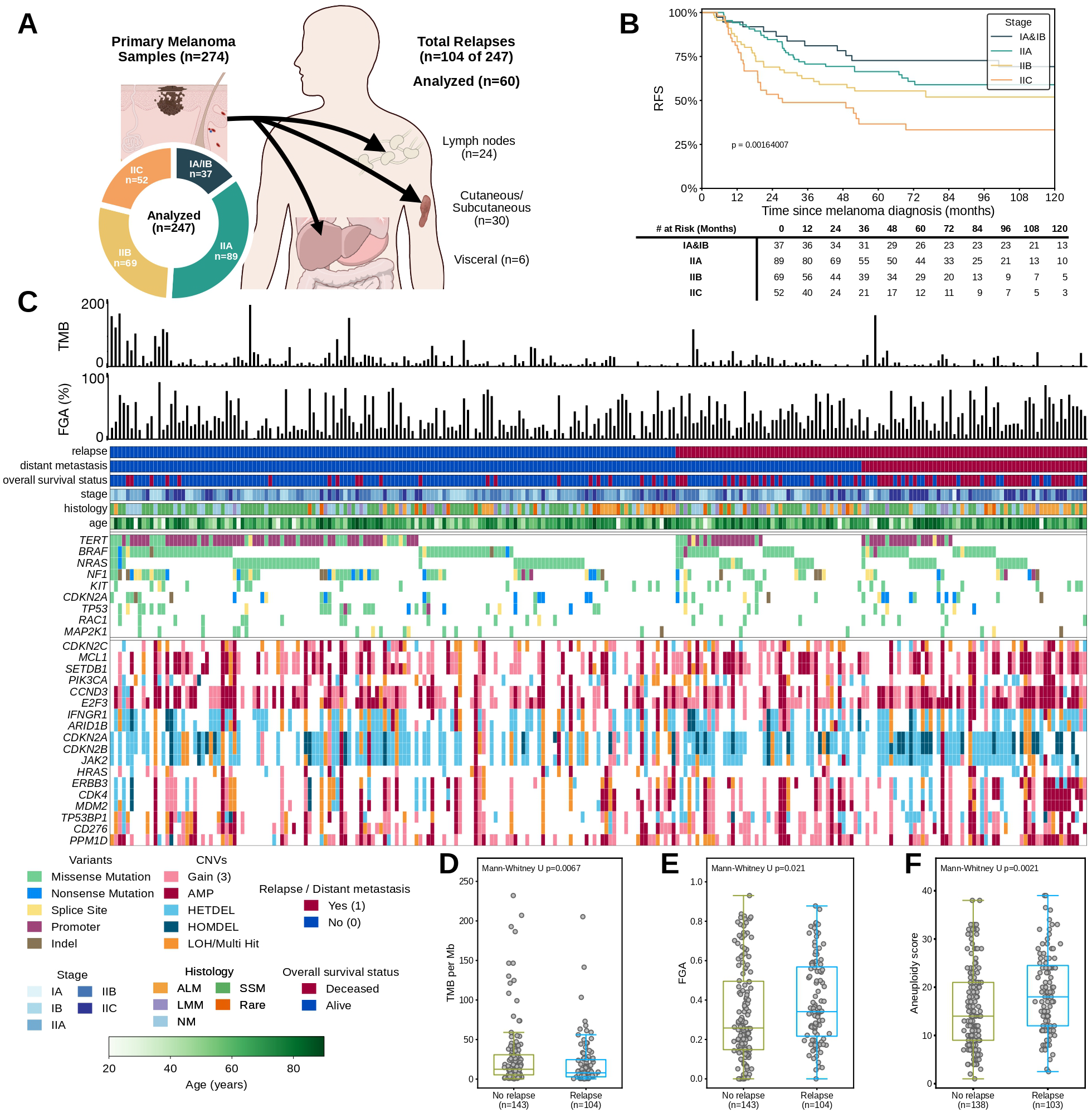
Clinical and genomic characteristics of primary cutaneous melanomas associated with metastatic relapse. A, Study design and cohort composition. Primary stage I/II cutaneous melanomas from 247 patients were characterized by targeted NGS. During follow-up, 104 patients developed metastatic relapse, and 60 corresponding metastatic specimens were analysed. B, Kaplan–Meier estimates of relapse-free survival (RFS) according to American Joint Committee on Cancer (AJCC) ninth-edition stage, with numbers at risk shown below (log-rank test, P = 0.0016). C, Integrated clinical and genomic landscape of the 247 primary melanomas, grouped by subsequent relapse status. Bar plots show the tumour mutational burden (TMB) and fraction of genome altered (FGA) for each tumour. Clinical annotation tracks indicate subsequent relapse, distant metastasis, overall survival status, tumour stage, histological subtype and age at diagnosis. The lower tracks show selected nucleotide variants and somatic copy-number alterations in major oncogenes and tumour-suppressor genes. D–F, Comparison of TMB (D; no relapse, n = 143; relapse, n = 104), FGA (E; n = 143 and n = 104, respectively) and aneuploidy score (F; n = 138 and n = 103, respectively) between primary tumours from patients who did or did not subsequently experience relapse. P values were determined using two-sided Mann–Whitney U-tests (D, P = 0.0067; E, P = 0.021; F, P = 0.0021). Points represent individual primary tumours; boxes indicate the median and interquartile range. ALM, acral lentiginous melanoma; AMP, amplification; HETDEL, heterozygous deletion; HOMDEL, homozygous deletion; LMM, lentigo maligna melanoma; LOH, loss of heterozygosity; NGS, nextgeneration sequencing; OS, overall survival; Rare, rare histological melanoma subtypes; SCNA, somatic copy-number alteration; SNV, single-nucleotide variant; SSM, superficial spreading melanoma.

Sentinel lymph-node biopsy was performed in 205/247 patients at diagnosis. 104 patients experienced metastatic relapse 44 developed loco-regional metastases, 34 loco-regional followed by subsequent distant metastases, 6 simultaneous loco-regional and distant metastases, and 20 upfront distant metastases. Median follow-up was 92 months (95% CI, 78–98); median relapse-free survival (RFS) was 152 months (95% CI, 69–not reached) and median distant-metastasis-free survival (DMFS) 217 months (95% CI, 168–not reached). As expected, increased tumour thickness and higher clinical stage were significant predictors of shorter RFS (e.g. thickness >4 mm: HR = 2.98, P < 0.001; Extended Data Fig. S 1A, Table S1).

All 247 primary CMs harboured multiple oncogenic mutations. *TERT* promoter mutations were the most frequent alteration (93/247; 37%). *BRAF*, *NRAS* and *NF1* driver mutations, which occurred in largely mutually exclusive patterns, were collectively detected in 179/247 tumours (72%), and *KIT* mutations in 16/247 (7%). Beyond these canonical drivers, recurrent copy-number alterations included amplifications of *CDK4*, *MCL1*, *CCND1* and *PIK3CA* and deletions affecting the *CDKN2A* and *CDKN2B* and *TP53BP1* tumour-suppressor loci (Fig. 1C). The median tumour mutational burden (TMB) was 9.25 variants/Mb (IQR, 3.19–22.16; Fig. 1D) and the median number of oncogenic mutations per tumour was 4 (IQR, 2–10; Extended Data Fig. S 1B). No individual mutation was associated with shortened RFS in univariate Cox analysis (Extended Data Fig. S 1C, Table S2). Most primaries also carried SCNAs, including amplifications of oncogene-containing regions and deletions of tumour-suppressor regions (Extended Data Fig. S 1D).

High TMB, which is associated with improved immunogenicity and better immunotherapy outcomes in metastatic melanoma, characterised primary tumours that did not relapse (P = 0.0067; Fig. 1D). Conversely, a higher fraction of genome altered (FGA)—a measure of chromosomal instability (P = 0.021; Fig. 1E)— and a higher aneuploidy score (P = 0.0021; Fig. 1F) were significantly associated with tumours that subsequently relapsed. Thus, global genomic instability, rather than any single-gene mutation, distinguished primary tumours destined for metastatic progression.

### An SCNA-based signature predicts relapse independently of clinical staging

In addition to global changes mediated by chromosomal instability, we asked whether prognostic information could be localised to specific SCNA loci. Amplifications at 1q21–q25, 12q12–q15 and 15q21– q26, and homozygous deletions at 9p21–p24 and 15q14–q15, were significantly enriched in tumours that later relapsed (Fig. 2A,B). Within these loci, our panel covered *MCL1, SETDB1* (1q21–q25), *CDK4, ERBB3* and *MDM2* (12q12–q15), *CD276* (15q24), *CDKN2A, CDKN2B* and *JAK2* (9p21–p24) and *TP53BP1* (15q15). From these genes we defined a copy-number signature termed *OncoCycle* of six genes from non-overlapping loci. Tumours were classified OncoCycle-positive if they carried an amplification of *CDK4*, *MCL1* or *CD276*, or a functional biallelic loss of *CDKN2A*, *CDKN2B* or *TP53BP1* (defined as homozygous deletion, or a loss-of-function oncogenic alteration with concurrent heterozygous deletion or loss of heterozygosity).

**Figure 2.**
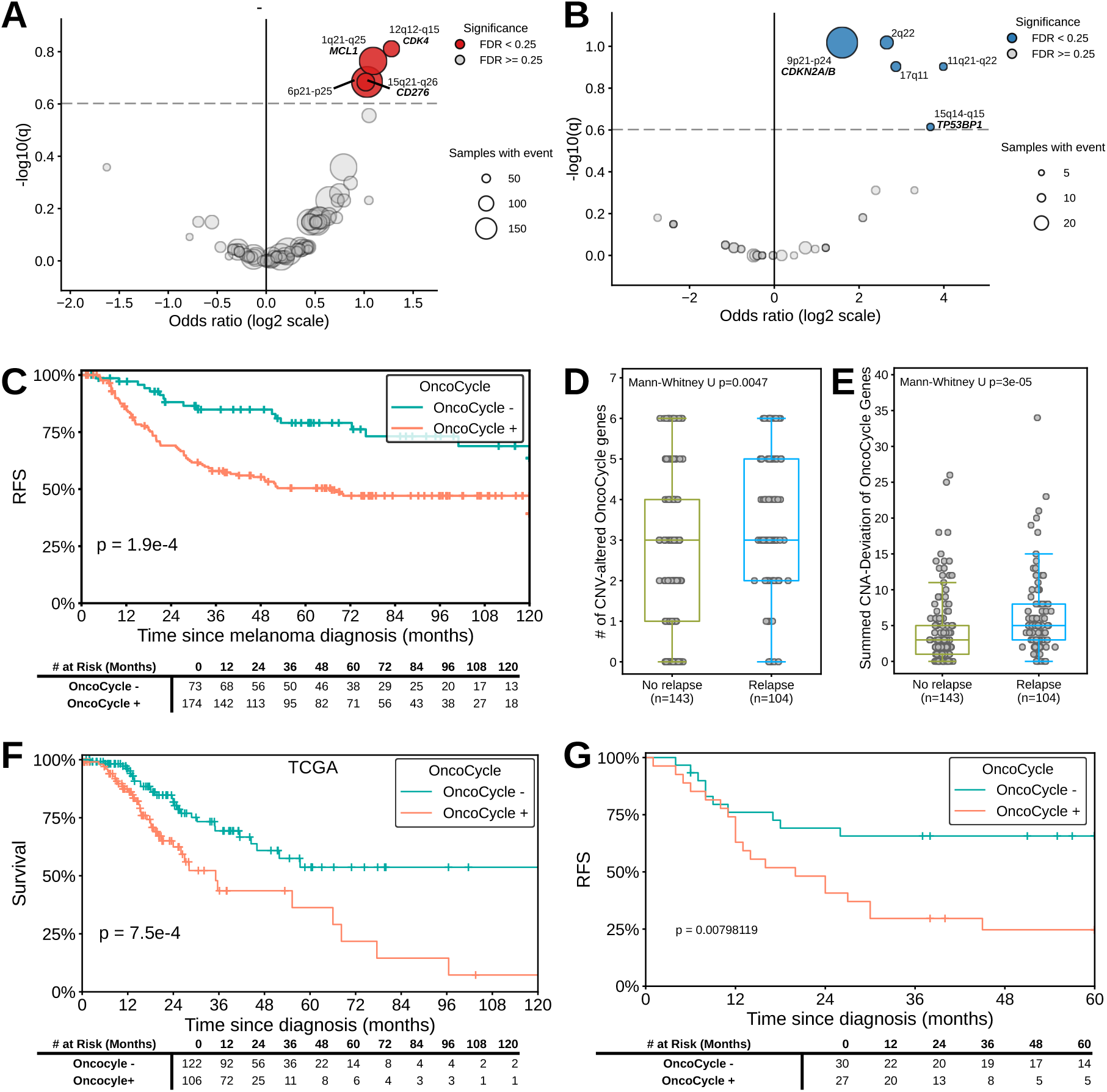
OncoCycle somatic copy-number alterations predict metastatic relapse and adverse survival. A,B, Association of focal amplifications (A) and homozygous deletions (B) in primary cutaneous melanomas with subsequent relapse. The x axis shows the log2-transformed odds ratio for relapse, and the y axis shows −log10(q), where q denotes the false discovery rate (FDR)-adjusted P value. The vertical line indicates an odds ratio of 1, the horizontal dashed line indicates an FDR threshold of 0.25, and circle size represents the number of primary tumours harbouring each event. Significant regions are coloured, and selected loci and candidate genes are annotated. C, Kaplan–Meier estimates of relapse-free survival (RFS) according to OncoCycle status in the discovery cohort (OncoCycle-negative, n = 73; OncoCycle-positive, n = 174; log-rank test, P = 2.5 × 10⁻⁴). Tumours were classified as OncoCycle-positive when they harboured amplification of *CDK4*, *MCL1* or *CD276*, or functional biallelic loss of *CDKN2A*, *CDKN2B* or *TP53BP1*. D,E, Comparison of the number of OncoCycle genes affected by copy-number alterations (D) and the cumulative magnitude of copy-number deviation from the diploid state across OncoCycle genes (E) between primary tumours from patients who did not relapse (n = 143) and those who relapsed (n = 104). P values were determined using the Mann–Whitney U-test (D, P = 0.0047; E, P = 3 × 10⁻⁵). Points represent individual primary tumours; boxes indicate the median and interquartile range. F, Kaplan–Meier estimates of overall survival from cBioPortal melanoma cohort sorted by OncoCycle status (OncoCycle-negative, n = 122; OncoCycle-positive, n = 106; log-rank test, P = 7.5 × 10⁻⁴). G, Kaplan–Meier estimates of RFS in an independent validation cohort (OncoCycle-negative, n = 30; OncoCycle-positive, n = 27; log-rank test, P = 8.0 × 10⁻³). Tick marks in C, F and G indicate censored observations, and numbers at risk are shown below each plot. CNV, copy-number variation; FDR, false discovery rate; RFS, relapse-free survival

Adding OncoCycle to AJCC 9^th^-edition staging^20^ significantly improved prognostic performance (likelihood-ratio test χ² = 12.39, df = 1, P = 4.3 × 10⁻⁴; n = 247, Table S3). In a fully adjusted model including AJCC9 stage, tumour thickness, ulceration, TMB and FGA, OncoCycle carried the highest hazard ratio (HR = 2.21, 95% CI 1.28–3.79, P = 0.004; Extended Data Fig. S 2A). OncoCycle-positive tumours had shorter RFS (P = 2.5 × 10⁻⁴; Fig. 2C, S2B) and DMFS (P = 0.037; Extended Data Fig. S 2C). Although positivity requires only a single alteration, relapsing tumours accumulated more altered OncoCycle genes (P = 0.0047; Fig. 2D) and a greater cumulative copy-number deviation from the diploid state across these genes (P = 3 × 10⁻⁵; Fig. 2E).

OncoCycle generalised beyond the discovery cohort. In The Cancer Genome Atlas^34,35^ via cBioPortal^36^, OncoCycle alterations predicted worse overall survival in primary melanoma and in primary lung, kidney, bladder, and pancreatic cancers (Fig. 2F, S2D). Furthermore, we performed whole-exome sequencing (WES) of 57 primary CM from an independent multicentre cohort with up to 60 months of follow-up, and found that OncoCycle positivity and stage (not shown) were significant predictors of RFS (P = 8.0 × 10⁻³; Fig. 2G). Thus, the dominant predictor of relapse was not oncogenic mutational status, but copy-number alteration concentrated at six loci.

### Copy-number evolution characterises metastatic progression

To resolve how individual tumours evolve during progression, we compared 60 matched primary– metastasis pairs, distinguishing within-patient changes from interpatient variation (Fig. 1A). TMB did not differ significantly between primaries and matched metastases (P = 0.064; Fig. 3A), indicating no net increase in mutational burden. In contrast, mutant-allele tumour heterogeneity (MATH) scores were significantly lower in metastases (P = 0.007; Fig. 3B), consistent with clonal selection during seeding, and FGA was significantly higher in metastases (P = 0.014; Fig. 3C), indicating chromosomal instability and increased copy-number burden. Metastases frequently harboured amplifications of *CDK4*, *CD276*, *MCL1*, *BRAF*, *NRAS* and *KIT* and losses of *CDKN2A/B*, *NF1*, *PTEN* and *TP53* (Fig. 3D).

**Figure 3.**
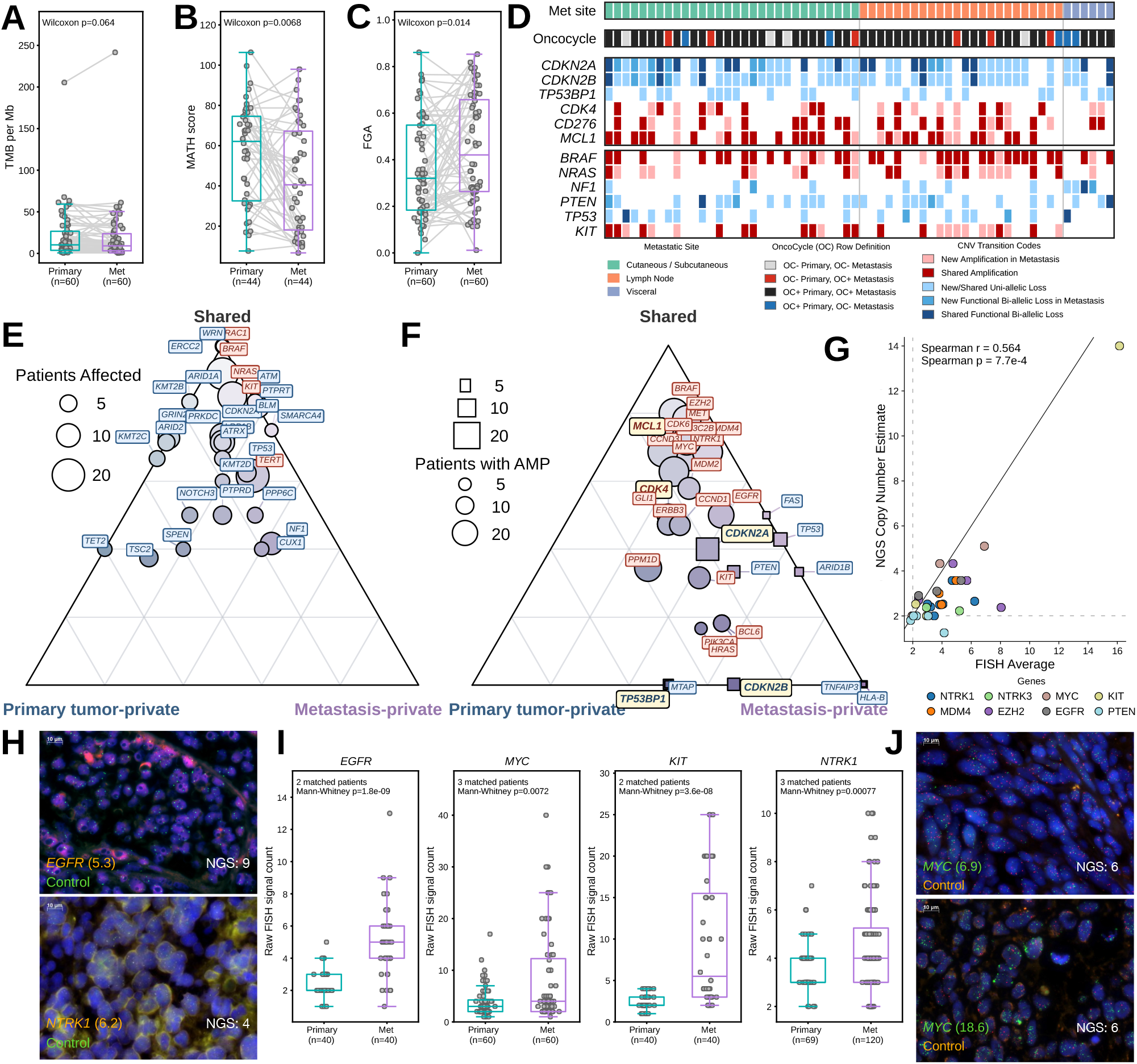
Somatic copy-number alterations define genomic evolution from primary to metastatic cutaneous melanoma. A–C, Paired comparison of TMB (A), MATH score (B) and FGA (C) between primary cutaneous melanomas and their matched relapses. Lines connect matched specimens. Primary tumours and relapses were compared using two-sided Wilcoxon signed-rank tests. Points represent individual tumours; boxes indicate the median and interquartile range. D, Copy-number changes in OncoCycle genes and other major oncogenes and tumour-suppressor genes across 60 matched primary– relapse pairs. Columns represent matched tumour pairs grouped by relapse site. The OncoCycle annotation indicates whether the primary tumour and matched relapses were OncoCycle-negative or OncoCycle-positive. Gene-level tracks show newly acquired or shared functional biallelic losses for tumour-suppressor genes and newly acquired or shared amplifications for oncogenes. E, Ternary plot showing the distribution of nucleotide variants that were shared between matched tumours, private to the primary tumour or private to the relapses. Circle size indicates the number of affected patients with said variant. F, Ternary plot showing the distribution of gene amplifications and functional biallelic losses that were shared, primary-tumour-private or metastasisprivate. Circles denote amplifications, squares denote functional biallelic losses and symbol size indicates the number of affected patients. G, Correlation between sample-average copy-number estimates derived from NGS and mean copy number measured by FISH for selected genes across 33 specimens. The solid line denotes the identity line, and dotted lines indicate a copy number of two representing a standard diploid. H, Representative FISH images of EGFR (orange), DAPI (green) and NTRK1 (orange), DAPI (green) in metastatic specimens. The mean FISH-derived copy number per nucleus is shown in parentheses, and the corresponding NGS-derived tumour copy-number estimate is indicated for each tumour. I, FISH signal counts for EGFR, MYC, KIT and NTRK1 in matched primary tumours and metastases. P values were determined using two-sided Mann–Whitney U-tests. Points represent individual nuclei; boxes indicate the median and interquartile range. J, Representative FISH images of a matched primary tumour and relapse of MYC (green), DAPI (orange). Target-gene and control probes are shown with nuclear counterstaining. NGS-derived tumour copy-number estimate is indicated when available. CNV, copy-number variation; FGA, fraction of genome altered; FISH, fluorescence in situ hybridization; LOF, loss of function; MATH, mutant-allele tumour heterogeneity; OC, OncoCycle; TMB, tumour mutational burden.

We next asked whether this copy-number divergence was accompanied by parallel mutation acquisition. Most oncogenic SNVs, including drivers in *BRAF*, *NRAS*, *NF1*, *TP53*, *KIT*, *ATM* and *ARID1A*, were shared between primary and metastasis, with few private variants (Fig. 3E). In contrast, SCNAs were disproportionately metastasis-specific, including amplifications of *MCL1*, *CCND1, CDK4*, *EZH2*, *MYC*, *PIK3C2B*, *NTRK1*, *MDM2*, *MDM4* and *EGFR* and biallelic losses of *CDKN2A*, *CDKN2B*, *PTEN* and *TP53* (Fig. 3F). Metastatic progression was therefore associated predominantly with increased SCNA burden rather than widespread acquisition of oncogenic mutations.

To validate the sequencing-derived copy-number estimates by an orthogonal method, we performed fluorescence in situ hybridisation (FISH) for nine genes across 33 samples (Fig. 3G–J). Total copy-number estimates from panel sequencing correlated with FISH (Spearman r = 0.51; Fig. 3G,H). FISH also confirmed significantly higher copy numbers of *EGFR*, *MYC*, *KIT* and *NTRK1* in metastases than in matched primaries (EGFR, P = 1.8 × 10⁻⁹; MYC, P = 0.0072; KIT, P = 3.6 × 10⁻⁸; NTRK1, P = 7.7 × 10⁻⁴; Fig. 3I). In patient FO13524, *MYC* copy number rose from 6 in the primary to 18 in the matched metastasis, and FISH confirmed the gain arose from chromosomal amplification (Fig. 3J).

### Clonal architecture of metastatic dissemination

The reduction in intratumoural heterogeneity suggested that progression involves the selective expansion of specific subpopulations within the primary tumour. We therefore asked whether metastases were seeded by a single dominant clone or by multiple subclones. Using PyClone to infer subclonal populations from variant allele frequencies, together with a custom clone-dominance metric based on the variants shared between each clone and the corresponding metastasis, we observed a wide distribution of dominance scores, providing evidence for both monoclonal and polyclonal seeding (Fig. 4A,B).

**Figure 4.**
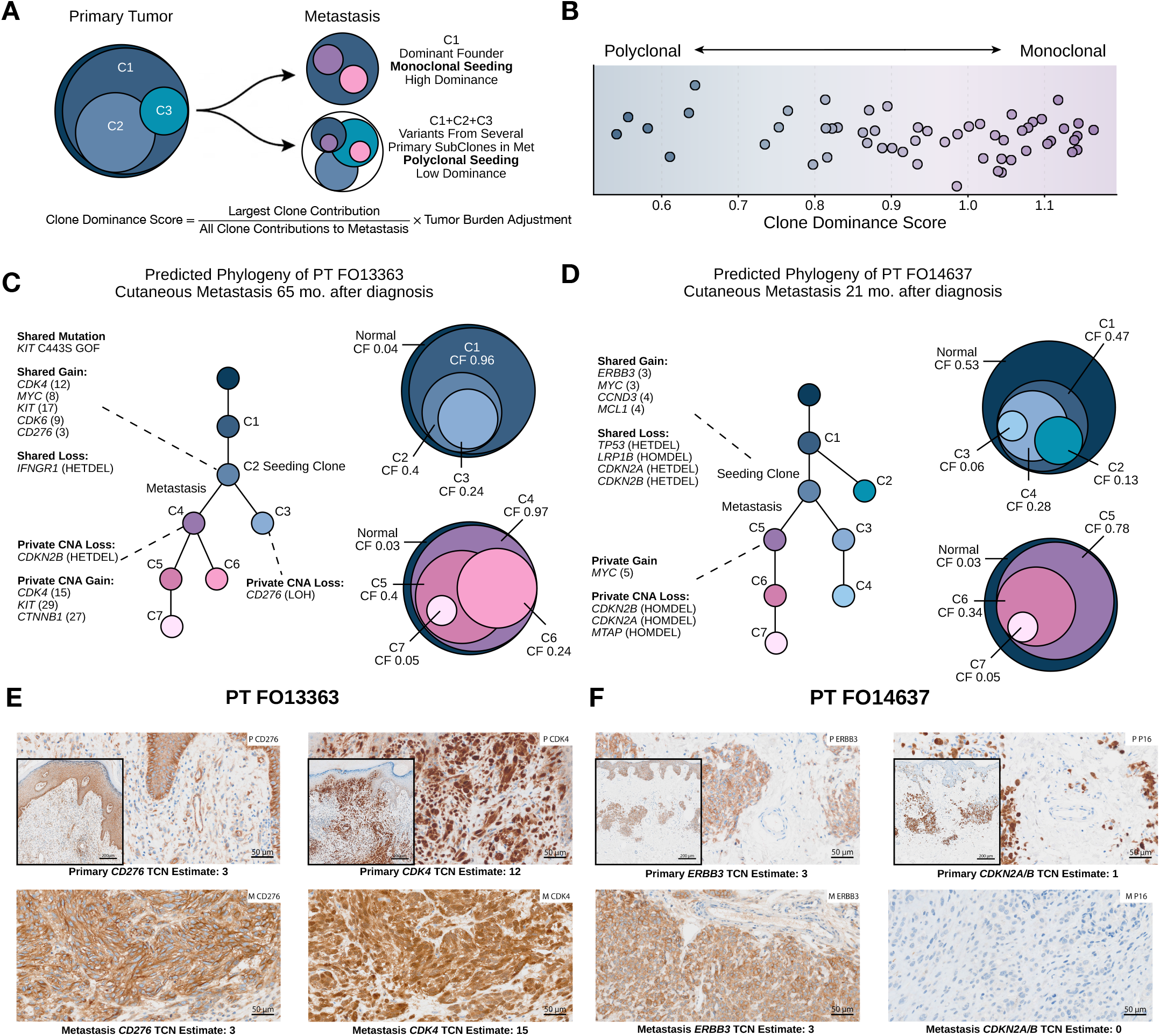
Clonal seeding from primary to metastatic melanoma. A, Schematic illustrating the calculation of the clone dominance score and its use in classifying matched primary tumour–metastasis pairs as arising through monoclonal or polyclonal seeding. The score was calculated as the contribution of the largest clone relative to the total clonal contribution to the metastasis, adjusted for tumour burden. B, Distribution of clone dominance scores across matched primary–metastasis pairs. Lower scores indicate more polyclonal seeding, whereas higher scores indicate greater dominance by a single metastatic founder clone. Each point represents one matched pair of a primary tumour and respective relapse. C,D, Reconstructed clonal phylogenies of two representative patients with cutaneous metastases: patient FO13363 (C), and patient FO14637 (D). Phylogenetic trees show the inferred relationships among clones, with the metastatic seeding clone indicated. Circle diagrams show the estimated clonal composition of the primary tumour and relapse; circle size represents clone fraction (CF) with adjustments for visualization purposes. Selected shared and metastasis-private mutations and copy-number alterations are shown, with copy-number estimates indicated in parentheses. E, Representative immunohistochemistry images from patient FO13363 showing CDK4 expression in the primary tumour and matched metastasis and the scattered CD276 expression in the primary and the strong CD276 expression in the metastasis. Corresponding NGS-derived total copy-number estimates for the altered tumour clone are shown. F, Representative immunohistochemistry images from patient FO14637 showing ERBB3 expression in the primary tumour and matched metastasis and the presence of p16 in the primary and loss of p16 expression in the metastasis, corresponding to the acquired homozygous deletion of *CDKN2A*. Corresponding NGS derived total copy-number estimates for the altered tumour clone are shown. Insets show higher-magnification views. CF, clone fraction; GOF, gain of function; IHC, immunohistochemistry; TCN, tumour copy number.

We reconstructed the clonal evolution of twelve and show two representative patients with monoclonal spread (Fig. 4C,D). In both, canonical driver mutations (e.g. *KIT*) and key OncoCycle SCNAs (e.g. *CDK4* or *MCL1* amplification) arose early during primary evolution and were fully preserved in the metastasisseeding clones. Following dissemination, these clones acquired additional, metastasis-specific SCNAs, including steep amplifications of *KIT*, *CDK4*, *CTNNB1*, *MDM4*, *SETDB1* or *MCL1* and biallelic loss of *CDKN2A*. Thus, OncoCycle SCNAs developed in the primary tumour before seeding, while subsequent subclonal evolution selectively enriched for further SCNAs that promote cell-cycle progression, chromosomal instability and immune evasive cues.

Immunohistochemistry (IHC) across 82 sections confirmed that these alterations are reflected at the protein level: structural amplifications correlated with strong protein expression, and biallelic deletions with complete loss of expression. In patient FO13363, a *CDK4* amplification present in both the primary (total copy number [TCN] = 12) and the metastasis (TCN = 15) corresponded to strong CDK4 staining in both. CNV calling identified a focal *CD276* amplification in a subpopulation of the primary tumour (0.1 < CCF < 0.3), which was markedly enriched in the metastasis (CCF > 0.8). This was consistent with the IHC, with focal areas of stronger B7-H3 staining in the primary tumour and homogeneous intense B7-H3 staining colocalizing with CDK4-positive cells in the metastasis (Fig. 4E). In patient FO14637, an *ERBB3* amplification shared by primary and metastasis was validated by robust HER3 staining, and a metastasisacquired homozygous deletion of *CDKN2A* corresponded to complete loss of p16 staining (Fig. 4F). These findings confirm that copy-number alterations retained or enriched during progression are expressed at the protein level and support the continued biological relevance of OncoCycle SCNAs as additional alterations accumulate.

### Spatial transcriptomics resolves clonal and microenvironmental evolution

We performed high-resolution spatial transcriptomic profiling of 19 primary CMs—8 from patients who subsequently relapsed and 11 from patients who did not—together with 6 matched relapses (Fig. 5A). Copynumber profiles inferred with inSituCNV resolved discrete tumour subclones and their spatial localisation within tissue architecture (Fig. 5B), recovering genomic alterations with concordance to DNA sequencing and strong sample-wise correlation across loss, neutral and gain states (Spearman ρ = 0.73; Fig. 5C).

**Figure 5.**
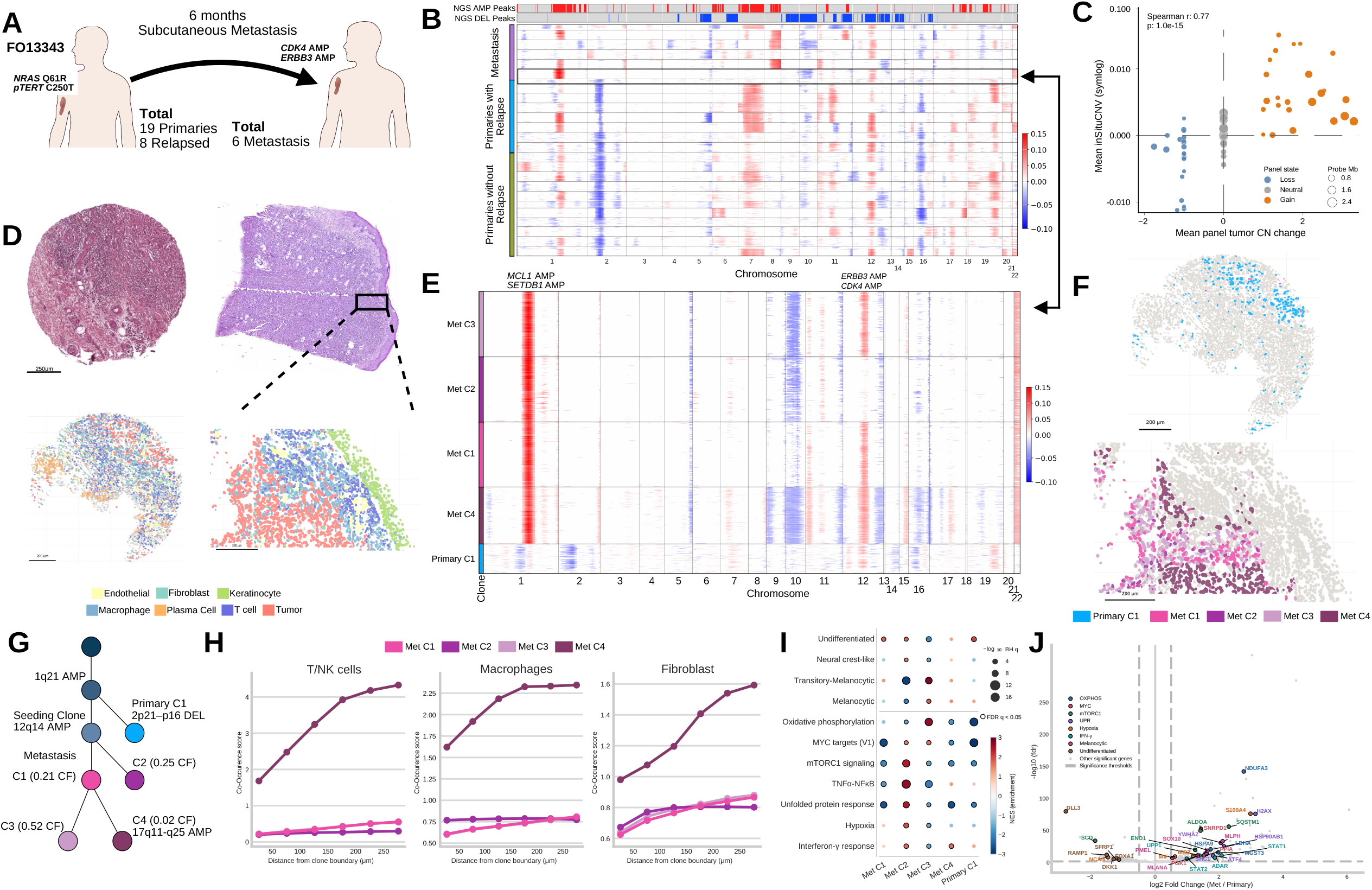
Spatial transcriptomics resolves clonal evolution and microenvironmental changes during metastatic progression. A, Study design of ST characterization of matched primary and metastatic melanomas. A representative primary tumour–metastasis pair from patient FO13343 and selected genomic alterations identified by NGS sequencing are indicated. B, inSituCNV-derived copy-number profiles across metastatic lesions, primary tumours from patients who relapsed and primary tumours from patients who did not relapse. Samples are ordered by clinical group. The upper tracks show recurrent amplification and deletion peaks identified by GISTIC^42^ analysis of panel-based NGS data for this cohort, aligned for comparison with the spatial transcriptomic copy-number profiles. C, Relationship between panel-based NGS copy-number states and sample-level inSituCNV measurements across all samples. Each point represents a sample–copy-numberstate aggregate (n = 72), with the mean NGS copy-number change shown on the x axis and the mean inSituCNV signal across genomic bins assigned to the same state within an individual sample shown on the y axis. Points are coloured according to the NGS-defined copy-number state (loss, neutral or gain) and scaled by the summed genomic span of the contributing bins. The y axis is displayed on a symmetric logarithmic scale (Spearman r = 0.729). D, H&E-stained sections, segmented cell transcriptome-based celltype assignments are shown for the paired primary tumour and metastasis, including an enlarged region of the metastatic specimen. E, Enlarged inSituCNV heatmap of the matched primary tumour and metastasis from patient FO13343, showing the copy-number profiles and subclonal distribution of the inferred tumour clones. F, Spatial distributions of inferred clones in the paired primary tumour and metastasis of patient FO13343, including an enlarged region of the metastatic specimen. G, inSituCNV-inferred phylogenetic relationship among primary and metastatic clones from patient FO13343. Nodes represent inferred clones, with clone fractions and selected accumulated copy-number alterations indicated. H, Spatial co-occurrence of T cells, macrophages and fibroblasts with individual metastatic tumour clones from FO13343 identified from ST. Co-occurrence scores are plotted as a function of distance from each clone boundary. I, Dot plot of gene-set enrichment for melanoma differentiation programs and selected cancer hallmark gene sets across the inferred metastatic tumour clones of patient FO13343. Dot colour indicated the normalized enrichment score (NES) and dot size scales with statistical significance (−log₁₀ FDR). Enrichment with FDR < 0.05 are outlined. J, Differential gene-expression analysis between primary and metastatic tumour populations from ST of patient FO13343. Selected significant genes are coloured according to the differentiation program and hallmarks. FDR, false discovery rate; H&E, haematoxylin and eosin; ST, spatial transcriptomics.

In patient FO13343, who developed a subcutaneous metastasis 6 months after primary tumour diagnosis, inSituCNV identified one major primary-tumour clone and four metastatic clones (Fig. 5D,E). We identified one spatially confined clone in the primary tumour that had shared truncal copy-number events in its corresponding metastatic lesion (Fig. 5F). This metastasis-competent clone had specific copy number gains spanning *MCL1* and *SETDB1* (1q21) and further diverged in the metastatic lesion, where it developed additional amplification in *ERBB3* and *CDK4* (12q13–q14) (Fig. 5E,G).

The four metastatic subclones occupied distinct cellular neighbourhoods (Fig 5F). Namely, Met clone 4 (Met C4) was strongly co-localized with T cells and macrophages, and modestly so with fibroblasts as compared to the other clones (Fig. 5H; Extended Data Fig. S 3a,b). In addition to their underlying SCNA patterns and associated TMEs, these subclones varied in their cancer cell intrinsic transcriptional programs. While the metastasis-competent clone in the primary tumour (Primary C1) was characterized by an undifferentiated melanoma lineage state, Met C3 in the metastasis strongly expressed the melanocytic-totransitory state and enriched for oxidative phosphorylation. Other clones, such as Met C1 maintained the undifferentiated state, and Met C2 co-expressed the neural-crest like stem-like state and a hypoxia program (Fig. 5I; Extended Data Fig. S 3c,d). In contrast, Met C4 did not show a strong bias towards a cell state along the melanoma differentiation spectrum. These cell state differences were accompanied by distinct pathway gene expression. Among others, Met C2 and Met C4 were enriched for interferon-gamma response pathway. This is consistent with Met C4’s lymphoid/macrophage rich TME and Met C2’s neural-crest like state, which has been associated with increased expression of antigen presentation machinery ^37^.

Consistent with this genomic and spatial divergence, primary and matched metastatic tumour cells displayed broad transcriptional differences (Fig. 5J). Cancer cells from metastatic lesions differentially expressed genes involved in oxidative phosphorylation (*e.g. LDHA*), MYC targets, mTORC1 signalling (*SQSTM1, ALDOA*), the unfolded-protein response (*HSP90AB1*) and hypoxia (Fig. 5J). Together, these analyses capture overall differences between primary tumour and metastasis cancer cells, but also highlight genomic and associated transcriptomic differences within the same lesion. Comparing the pooled transcriptomics data of the 19 primaries with those of the 6 metastatic melanomas provided equivalent results (Extended Data Fig. S 3e). Among the genes higher in metastases were several associated with migration and invasion, including *PTK2* (focal adhesion kinase), *LGALS3* (galectin-3), *NES* (nestin) and *VIM* and in particular, the metastasis-associated *S100* protein *S100A4.* The glycolytic and hypoxia-responsive genes *PGK1, NDRG1* and *NAMPT* were also higher.

## Discussion

By pairing integrating copy-number, mutational, and spatial analyses of primary melanomas with treatment-naïve, matched metastases and long-term clinical follow-up, we define the evolutionary trajectory at unprecedented depth and resolution. Important conclusions emerge from our work. First, the alterations that distinguish tumours destined to metastasise, and that accumulate during metastatic progression, are SCNAs rather than point mutations. Second, metastatic competence is often established early, encoded by a chromosomal instability and a compact copy-number programme present in the primary tumour, and then elaborated by continued accumulation of SCNAs in genes coding for cell cycle control, chromosomal stability or immune clearance during dissemination.

These findings contribute to resolving a long-standing paradox. Although metastasis is widely assumed to require a heritable pro-metastatic trait, metastasis-specific driver mutations have been difficult to identify, and oncogenic mutations are largely shared between primary tumours and their metastases^1^. Our data are consistent with this, showing that oncogenic SNVs, including the canonical melanoma drivers (explaining the efficacy of adjuvant BRAF inhibition)^33^, were shared between compartments. In contrast, SCNAs dominated the metastasis-specific alterations. They accounted for both, the molecular identity of the heritable trait, and the genomic divergence between primary and metastasis. SCNAs, on a background of chromosomal instability (CIN), thus provide the heritable, selectable substrate for metastatic evolution that recurrent point mutation does not. While CIN is now a well-established hallmark of metastatic lesions in melanoma and other cancers^13,14,22,25,28,38^ little is known about how CIN in primary tumours dictates metastasis risk and progresses in matched lesions.

Whether metastatic capacity is intrinsic to a tumour from its inception or acquired stepwise during growth has been debated for decades^1–3^. Our findings suggest these views are not mutually exclusive in melanoma. The OncoCycle programme is already present in the primary tumour, predicts relapse independently of stage, and, in monoclonal pairs, is carried by the clone that seeds the metastasis, consistent with an early, largely intrinsic determinant of metastatic competence that is detectable in bulk tissue^39^. At the same time, disseminated clones continue to acquire SCNAs (for example *MYC*, *EGFR*, *NTRK1* and *MDM2*/*MDM4*, alongside further cell-cycle and interferon-pathway alterations) while intratumoural heterogeneity declines, consistent with ongoing, selection-shaped evolution after seeding. Metastatic competence in melanoma is therefore neither wholly fixed nor wholly acquired. Instead, tumours “born-to-be-bad” may still require additional driver events in the form of SCNAs to engender metastatic competence. The critical role of the OncoCycle programme in the metastatic progression is strongly supported by experimental data in mice, where genomic loss of *CDKN2A* or activating *CDK4*-mutations transform being melanocytic tumours into tumours prone to develop metastases^29–31^.

Clonal reconstruction further indicates that melanoma does not follow a single mode of dissemination. Clone-dominance analysis of matched pairs revealed both monoclonal and polyclonal seeding, and spatial transcriptomics resolved multiple copy-number-defined metastatic subclones derived from a shared seeding clone within an individual lesion. The accompanying reduction in heterogeneity from primary to metastasis is consistent with an evolutionary bottleneck at seeding, after which chromosomal instability regenerates subclonal diversity at the metastatic site.

Spatial profiling begins to connect these genomic events to their microenvironmental context. Copynumber-defined metastatic subclones occupied distinct immune and stromal niches, with one clone enriched at an immune-rich boundary, suggesting that subclonal genotype and local immune and stromal composition are coupled. This is biologically coherent given the OncoCycle genes themselves, including amplification of immunoinhibitory *CD276* (B7-H3) and recurrent alteration of interferon-pathway and cellcycle regulators provide plausible routes to immune evasion alongside a proliferative advantage^40^.

Clinically, OncoCycle converts these observations into a prognostic tool. It significantly improved on AJCC9 staging in treatment-naïve stage I/II disease, carried the highest hazard ratio in a fully adjusted model, and predicted outcome across several additional cancer types, indicating that the underlying copynumber programme is not melanoma-specific^41^. Because the cohort predates efficacious adjuvant therapy for early-stage melanoma, OncoCycle captures the natural history of metastatic risk and is well suited to identifying early-stage patients who might benefit from adjuvant treatment.

Our study has limitations. Copy number was assessed primarily by targeted panel sequencing rather than multi-region whole-genome sequencing, which limits lineage resolution. We mitigated this with orthogonal FISH, protein validation and single-cell spatial copy-number inference, and a multicentre validation cohort where CNAs were assessed by WES. Furthermore, we have previously demonstrated that copy number inferences at gene and chromosome arm level can be robustly determined from panel sequencing data^26^. However, our clone-dominance metric should be interpreted as an approximation rather than a formal measure of clonal diversity. Most matched pairs comprised a single, metachronous metastasis, so late seeding and metastasis-to-metastasis spread cannot be fully excluded, and we did not formally quantify the randomness of metastatic seeding. While our single-cell spatial transcriptomics cohort represents among the largest in the melanoma field, continued expansion of such efforts are needed to further examine our initial findings.

Overall, our data support a model in which focal CNAs on a background of chromosomal instability prime metastatic competence early in primary melanoma and continue to evolve during dissemination. They thus offer a mechanistic account of melanoma progression, a clinically actionable measure of metastatic risk, 312 and identify potential targets for adjuvant therapies.

## Acknowledgements

This work was supported by Wilhelm Sander-Stiftung (2020.100.1 and 2025.216.1). Cluster of Excellence iFIT (EXC 2180) “Image-Guided and Functionally Instructed Tumor Therapies”, University of Tü bingen, Germany, funded by the Deutsche Forschungsgemeinschaft (DFG, German Research Foundation) under Germany’s Excellence Strategy -EXC 2180 – 390900677 (to M.R.), the National Institutes of Health (NIH) through National Cancer Institute (NCI) grants R01CA266446, R01CA280414, R37CA258829, and U54CA274506 (to B.I.), a Velocity Fellows Award, the Louis V. Gerstner, Jr. Scholars Program, a Tara Miller Young Investigator Award by the Melanoma Research Alliance (MRA), a Tara Miller Team Science Award for Metastasis Research by the MRA, a Leveraged Finance Fights Melanoma (LFFM) – MRA Team Science Award, and the Pershing Square Sohn Cancer Research Alliance Award (to B.I.). B.I. is a Cancer Research Institute Lloyd J. Old STAR (CRI5579). This work was additionally supported by the Herbert Irving Comprehensive Cancer Center (HICCC) (P30CA013696) Human Tissue Immunology and Immunotherapy Initiative (to B.I.). OR was supported by a structural grant from the DFG (German NGS competence center) and the EU (JA.PCM), S.N and J.S. by the Deutsche Forschungsgemeinschaft under the German National Research Infrastructure for Immunology (NFDI4Immuno) [NFDI 49/1 - 501875662], and NFDI 1/1 “GHGA - German Human Genome-Phenome Archive” (#441914366 to S.N.). S.N. was supported by the Deutsche Forschungsgemeinschaft under Germany’s Excellence Strategy (Grant EXC2180-390900677) and the Carl Zeiss Foundation “Certification and Foundations of Safe Machine Learning Systems in Healthcare TS, TA, were supported by the Cluster of Excellence iFIT “Image-Guided and Functionally Instructed Tumor Therapies” (EXC 2180/10072-1_1 and S.54.10072), University of Tübingen, Germany, funded by the Deutsche Forschungsgemeinschaft (DFG, German Research Foundation) under Germany’s Excellence Strategy-EXC 2180—390900677. TA, and TS received research funding from Novartis Pharma GmbH (S2TAF-057). We thank Professor Dr. Claudia Günther for her continued support, and Profs. Dr. M. Berneburg, J. Bauer and A. Yazdi for the multicenter analysis. We thank all colleagues who worked with the patients, and all patients who consented and motivated us to perform this analysis.

## Conflict of interest disclosures

B.I. has received consulting fees/honoraria from Volastra Therapeutics Inc, Merck, AstraZeneca, Novartis, Eisai, and Janssen Pharmaceuticals and has received research funding to Columbia University from Alkermes, Arcus Biosciences, Checkmate Pharmaceuticals, Compugen, Ideaya Biosciences, Immunocore, Merck, Regeneron, and Synthekine. B.I. is a scientific founder of Basima Therapeutics, Inc. IB received speaker fees from Novartis, Bayer and AstraZeneca and honoraria for advisory board participation from BMS and Novartis outside the submitted work. S.A.E, received funding from Almirall S.A. and BMS, and declares the pending European patent applications: OncoCycle (EP application no. 24 210 236.6), and OncoCycle (EP application no. 225 221 449.9). T.S. reports institutional funding from Novartis and Pierre-Fabre outside and the pending European patent applications: OncoCycle (EP application no. 24 210 236.6). L.F. reports research grants from Hookipa Pharma, SAKK/Immunophotonics, DFG Grant (Deutsche Forschungsgemeinschaft), Deutsche Krebshilfe, Philogen and Mundipharma; consulting fees from Philogen; participation on Data Safety Board University of Basel. C.S. reports an grant by Illumina, Inc., funding from BMS Stiftung Immuntherapie and Westdeutsche Studiengruppe. S.F. received personal honoraria from Kyowa Kirin, Stemline, Recordati Rare Diseases (speaker’s honoraria, advisory board), and grants from BioNTech SE, Neracare, SkylineDX and the EU MELCAYA project. C.G. reports fees for advisory board membership from MSD and Philogen; from CeCaVa and NeraCare; fees for expert testimony from NeraCare; and a nonremunerated role as President of the European Association of Dermato-Oncology (EADO). S.O. received travel support and speaker fees from Illumina, Inc., Oxford Nanopore Technologies. T.A. reports fees for advisory board membership from Delcath and Philogen; fees as invited speaker from Bristol Myers Squibb (BMS), NeraCare, Novartis and Pierre Fabre; fees for writing from CeCaVa and Medtrix; principal investigator for Agenus Inc., AstraZeneca, BioNTech, BMS, HUYA Bioscience, Immunocore, IO Biotech, Merck Sharp & Dohme (MSD), Pfizer, Philogen, Regeneron, Roche and University Hospital Essen; institutional fees as coordinating PI from Unicancer; institutional research grants from iFIT and Novartis; institutional funding from MNI Naturwissenschaftliches und Medizinisches Institut, NeraCare, Novartis, Pascoe, Sanofi and Skyline-Dx; membership of the American Society of Clinical Oncology (ASCO) and the Portuguese Society for Medical Oncology; and clinical expert for medical oncology for Infarmed. C.M.S. is a cofounder, shareholder and employee of Vicinity Bio GmbH, and scientific advisor to Enable Medicine Inc.. U.L. declares support from Merck Sharp & Dohme and the Deutsche Forschungsgemeinschaft (DFG), speakers and advisory board honoraria from Bristol Myers Squibb, Merck Sharp & Dohme, Sun Pharma, Sanofi, Almirall, Incyte, Skyline DX, Regeneron and Novartis; meeting support from Pierre Fabre, and Sun Pharma. A.J.S. reports consulting, funding, advisory, or lecture fees from Regeneron Pharmaceuticals Inc., Replimune Ltd; with Merck & Co Inc; Genesis Pharma; L’ Oreal. O.R. received funding from the German Research Foundation for the NGS Competence center and the EU funded project JA.PCM, grants from the EU, the BMFTR, Illumina and ONT and declares the following pending European patent applications: CIS-RESIST (EP application no. EP4397776A3), OncoCycle (EP application no. 24 210 236.6), and OncoCycle2 (EP application no. 225 221 449.9). F.F. has received speaker homoraria from Astra Zeneca, Aether AI, Stemline, Roche and Thermo Fisher and research support from Stemline and Thermo Fisher. M.R.reports grants for clinical studies from Agenus, Alcedis, Almirall Hermal, AstraZeneca, Bion-tech, Bristol-Myers Squibb, COLDPLASMATECH, Genentech, Horizon Therapeutics, HUYA Bioscience, HY-Pharm, Immunocore, Incyte, InflaRx, IO Biotech, IQVIA, Kartos Therapeutics, Leo Pharma, MSD Sharp & Dohme, Mühlenkreiskliniken Minden, NeraCare, Novartis Pharmaceuticals, Pfizer, Philogen, Pierre Fabre, Regeneron Pharmaceuticals, Replimune Group, RHEACELL, Roche, Sanofi-Aventis, SkylineDx, Sun Pharma, Takeda Pharmaceutical, Technische Universität Dresden, Unicancer, Universitätsklinikum Essen, Universitätsklinikum Freiburg, Universitätsklinikum Heidelberg, and grants for research projects from Deutsche Forschungsgemeinschaft, Deutsche Krebshilfe and Wilhelm Sander-Stiftung and the pending European patent applications: CIS-RESIST (EP application no. EP4397776A3), OncoCycle (EP application no. 24 210 236.6) and OncoCycle2 (EP application no. 225 221 449.9). E. C. declares the following pending European patent applications: OncoCycle (EP application no. 24 210 236.6), and OncoCycle2 (EP application no. 225 221 449.9). S.H. received personal honoraria from BMS, MSD, Regeneron & Pierre Fabre (speaker’s honoraria, advisory board) outside the submitted work.K.L., P.S, F.S, J.A., A.B.M, J.S, E.M, S.M, Y.M.H., L.S., O.T.P, M.K, S.H, P.M, S.N., declare no conflicts of interest.

## Figure legends

**Extended Data Figure 1: Clinical and genomic features associated with metastatic relapse in primary cutaneous melanoma.**

A, Univariate Cox proportional-hazards analysis of clinical features associated with relapse-free survival (RFS). Points indicate log hazard ratios (HRs) and horizontal bars indicate 95% confidence intervals; the vertical dashed line denotes *log* HR = 1. B, Number of oncogenic mutations per primary melanoma in patients who did not subsequently relapse (n = 143) and those who relapsed (n = 104). P value was determined using a two-sided Mann–Whitney U-test (P = 0.25). Points represent individual primary tumours; boxes indicate the median and interquartile range. C, Univariate Cox proportional-hazards analysis of recurrent oncogenic mutations and RFS. The number of tumours harbouring each alteration is indicated. Points indicate HRs and horizontal bars indicate 95% confidence intervals; the vertical dashed line denotes HR = 1. D, Genome-wide GISTIC2 profiles of recurrent amplifications (red) and deletions (blue) in primary melanomas from patients who did not subsequently relapse (left) and those who relapsed (right). Chromosomes are arranged vertically and recurrently altered cytobands are indicated.

AJCC9, American Joint Committee on Cancer ninth edition; HR, hazard ratio; SLNB, sentinel lymph-node biopsy.

**Extended Data Figure 2: OncoCycle predicts adverse outcomes independently of clinical features and across cancer types.**

A, Multivariable Cox *log* proportional-hazards analysis of RFS incorporating OncoCycle status, AJCC9 stage, tumour thickness, ulceration, TMB, FGA. B, Kaplan–Meier estimates of RFS jointly stratified by AJCC9 stage and OncoCycle status. Numbers at risk are shown below the plot; the overall P value was determined by log-rank test. C, Kaplan–Meier estimates of distant-metastasis-free survival (DMFS) according to OncoCycle status in the discovery cohort (OncoCycle-negative, n = 73; OncoCycle-positive, n = 174; log-rank P = 0.037). D, Kaplan–Meier estimates of overall survival in primary lung, bladder, pancreatic, and kidney cancers from cBioPortal, stratified by OncoCycle status. P values shown were determined using log-rank tests. Tick marks in B–D indicate censored observations.

**Extended Data Figure 3: Tumour-clone microenvironment composition and hypoxia gradients relative to vasculature in metastatic melanoma**

A, Non-tumour neighbour composition within 50 µm of each metastatic tumour clone. B, Neighbourhood enrichment analysis of spatial co-localization for each metastatic clone (Met C1–C4). Cell colours indicate the enrichment z-score for each cell type within a 50 µm neighbourhood; asterisks denote significance at FDR q < 0.05. C, Representative spatial section of the tumour clonal distribution (left) alongside the matched tumour cells coloured by per-cell HALLMARK_HYPOXIA score. Endothelial cells are highlighted in green. D, Median HALLMARK_HYPOXIA score of tumour cells as a function of binned distance to the nearest endothelial cell. E, Differential gene-expression analysis between primary and metastatic tumour populations from ST of 19 primary and 6 metastatic patients. Log_10_FDR (Y-axis) values were capped at 20 and values above 20 are plotted at triangles.

## Methods Ethical approval

This study was approved by the ethics committees of the institutions involved and was conducted in accordance with consensus ethical principles derived from international ethical guidelines, including the Declaration of Helsinki. This study was approved by the Ethics Commission of the Eberhard Karls University Tuebingen with the number 883/2019BO2.

## Clinical cohort

Here we analysed the genomes of primary cutaneous melanomas, healthy tissue and, where applicable, metastases from 271 patients diagnosed with stage IA to IIC cutaneous melanoma between 2000 and 2018. Patients were prospectively followed by the Central Malignant Melanoma Registry (CMMR) and received treatment at the Department of Dermatology of the University in Tuebingen, Germany^43^. In 247 patients we successfully sequenced both primary melanomas and healthy tissue. To be included in the study, patients had to meet the following criteria: (1) age ≥18 years, (2) a confirmed diagnosis of Primary Cutaneous Melanoma by a certified dermato-pathologist, and (3) no known germline mutations. Demographic information, clinicopathological characteristics, and survival data of the patients were extracted from the CMMR. Staging was conducted in accordance with the American Joint Committee on Cancer (AJCC) ninth Edition guidelines^20^. We obtained primary melanoma and normal tissue and where available a matched metastasis from archived Formalin-Fixed Paraffin-Embedded (FFPE) specimens. Normal skin or tumour-free sentinel lymph nodes served as normal tissue. New Hematoxylin and Eosin (HE) sections were prepared, and a certified dermatopathologist reviewed the diagnosis of primary cutaneous melanoma. Tumour content was quantified at the macroscopic level using HE-stained slides, and only samples with more than 5% tumour cells were included. Samples with a concentration of less than 4 ng DNA/ml were excluded. As confirmation cohort, we included 57 patients with stage I/II primary melanoma from five different centers.

## Sample preparation Sequencing

Next-generation sequencing (NGS) of both tumour and normal tissue was performed at the Institute of Medical Genetics and Applied Genomics, Medical Faculty Tuebingen^44^. Regions of interest were selectively enriched using the SureSelect XT Low Input Target Enrichment System from Agilent Technologies, Santa Clara, CA, USA. Hybrid capture was performed with custom-designed bait sets from SureSelect Somatic Cancer Panel v4, covering 396 cancer-related genes, and Sure Select Somatic Cancer Panel v5, covering 708 cancer-related genes, seven promoter regions, and selected intronic regions involved in gene fusions. 195 paired samples sequenced with v4 and 71 with v5. For WES libraries were processed using the Library Preparation EF Kit v2 with a custom exome enrichment probe kit (Twist Bioscience). Libraries were sequenced on an Illumina platform (NovaSeq6000 or NovaSeqXPlus). Quality control parameters, like sample or data swaps as well as all meta data were collected during all analysis steps

## Multiplex immunohistochemistry and FISH

Immunohistochemical studies were performed on 2-3 µm FFPE sections using the Ventana Ultra automated staining system (Ventana Medical Systems, Tucson, AZ, USA) and Ventana reagents, following the manufacturer’s protocols. The antibody panel included: p16 (clone INK4a, ready-to-use; Roche Diagnostics, Cat. No. 06594441001), with CC1 for 64 minutes; CDK4 (clone EPR4513, dilution 1:750; Abcam, Cat. No. ab108357), CC1 for 64 minutes; MDM2 (clone 3G187, dilution 1:25; Zytomed, Cat. No. 113-0230), CC1 for 40 minutes; HER3/ErbB3 (clone EBR22669-25, dilution 1:100; Abcam, Cat. No. ab256504), CC2 for 60 minutes; CD276/B7-H3 (clone BSB2813, dilution 1:75; Medac, Cat. No. RBT-B7H3), CC1 for 64 minutes; MYC (clone Y69, ready-to-use; Roche Diagnostics, Cat. No. 06504612001), CC1 for 64 minutes; and Cyclin D3 (CCND3; clone DCS22, dilution 1:400; Cell Signaling Technology, Cat. No. 2936). Staining intensity and percentage of positive cells was semiquantitatively assessed, and intensity was scored as negative (0), weak (1), moderate (2), or strong (3).

Fluorescence in situ hybridization (FISH) was performed according to the manufacturers’ instructions (Abbott Molecular, Zytomed Systems). In brief, 3.5-µm sections of formalin-fixed, paraffin-embedded (FFPE) tissue were deparaffinized in xylene (3 × 10 min, room temperature), dehydrated in 100% ethanol (2 × 5 min, room temperature), and air-dried for 5 min at room temperature.

Deproteinization was achieved by incubation in sodium citrate buffer (2× SSC, pH 6.0, 40 min, 95°C), followed by pepsin digestion (250 µg/mL in 0.9% sodium chloride, pH 2.0, 4 min, 37°C). The sections were subsequently dehydrated in ethanol (70% for 2 min, followed by 100% for 2 min, room temperature) and air-dried.

The FISH probes were applied to the sections, which were then covered with glass coverslips and sealed with Fixogum (Marabu, Germany). Probe and target DNA were co-denatured in a hybridization oven (Hybrite, Abbott Molecular) for 10 min at 75°C, followed by overnight hybridization at 37°C. After hybridization, the coverslips were removed, and post-hybridization washes were performed according to the respective probe manufacturers. For Abbott Molecular probes, the sections were washed in 2× SSC for 10 min at room temperature, followed by 2× SSC/0.3% NP-40 for 5 min at 73°C and a final wash in 2× SSC for 10 min at room temperature. For Zytomed Systems probes, the sections were washed three times in Wash Buffer A for 5 min each at 37°C. For the homebrew probe, the sections were washed in 2× SSC for 6 min at room temperature, followed by 2× SSC for 6 min at 76°C and 0.4× SSC for 6 min at room temperature. Subsequently, the sections were incubated with CAS-Block (Invitrogen) for 10 min at room temperature, followed by incubation with a fluorescently labelled anti-streptavidin antibody (1:200 dilution in CAS-Block; Invitrogen) for 60 min at room temperature. Finally, the sections were washed in 0.4× SSC for 6 min at room temperature.

After completion of the post-hybridization washes, all sections were dehydrated in graded ethanol (70%, 85%, and 100%; 1 min each, room temperature). DAPI (Abbott Molecular or Zytomed Systems) was applied, and the sections were coverslipped with glass coverslips.

## Bioinformatics analysis Processing of sequencing data and Alignment of reads

For samples with multiple sequencing runs or fragmented files, raw reads were concatenated by mate direction before alignment. We used an unmapped BAM (uBAM)-to-mapped BAM workflow following Genome Analysis Toolkit (GATK) best practices^45^. Raw paired-end FASTQ files were converted to uBAM format using FastqToSam. The uBAM files were subsequently standardized using RevertSam. Illumina adapter sequences were identified and tagged using MarkIlluminaAdapters before alignment. All GATK tools were run using GATK v4.6.2.0. Samples were excluded if more than 5% of reads lacked a mate, fewer than 90% of reads aligned to the reference genome, mean on-target coverage was below 100× for tumour samples or 50× for matched-normal samples.

Alignment to the GRCh38 human reference genome was performed using BWA-MEM v0.7.18^46^. SamToFastq extracted interleaved reads from the adapter-tagged uBAM files. BWA-MEM performed Smith-Waterman extension to generate base-level alignments. The mapped alignments were then piped into MergeBamAlignment. MarkDuplicates identified and flagged PCR and optical duplicates.

Coordinatesorted BAM files and their corresponding indices were generated and validated using SAMtools v1.21^47^.

## Calling somatic mutations

Somatic variants were called and filtered using GATK v4.6.2.0. Mutect2 was run in tumour-normal mode to generate unfiltered somatic variant calls, and read-orientation artifacts were modelled using LearnReadOrientationModel^48^. GetPileupSummaries was run separately on the tumour and matchednormal alignment files, restricted to the targeted panel intervals, using common single-nucleotide polymorphisms from gnomAD v3.1.1 as the population resource^49^. CalculateContamination was then used to estimate cross-sample contamination. FilterMutectCalls was applied to the unfiltered Mutect2 output.

## Variant Annotation

Annotations were provided by SnpEff version 5.3a, database GRCh38.86^50^. Upstream or regulatory variants were reclassified as promoter events if within 250 bp of a gene (300 bp for TERT). Coding sequence variants were retained only if an amino acid change was present, and frameshifts were classified as insertions or deletions based on allele length differences. Variants were deduplicated per gene and sample. If multiple alteration classes occurred within the same gene for a single sample, only the most severe class was retained, rather than labelling the sample as multi-hit for Figure 1C.

A second annotation set was generated with OncoKB v3.4.1-16^51^. Variants with a tumour VAF below 0.05 were excluded. The retained alleles were exported as minimal genomic-change MAF records and annotated with the OncoKB MafAnnotator against GRCh38 using the melanoma (CM) tumour type.

## Copy-number variant computation

Genomic intervals were partitioned into on-target and off-target regions. On-target regions were defined using the panel capture BED files. Off-target regions were represented by 100-kb genomic bins after exclusion of targeted intervals. Bins smaller than 50 kb were removed. Tumour and normal read-depth matrices were generated across these regions using BedCoverage. Off-target coverage 668 required a minimum mapping quality of 10 reads to restrict analysis to more confidently aligned background reads.

To incorporate allelic imbalance into copy-number calling, B-allele frequencies were used. Analysis was restricted to biallelic single-nucleotide polymorphisms with non-missing genotypes, valid allelic-depth values, and a total read depth of at least 10. At each locus, the B-allele frequency was defined as the fraction of reads supporting the alternate allele. When the same sample appeared in more than one overlapping pair, the observation with the greatest read depth was retained.

Copy-number analysis was performed separately within panel-specific sample subsets using ClinCNV v1.19.152. Normalized on-target and off-target coverage matrices were analysed together with the filtered B-allele-frequency profiles. A minimum segment length of three bins and a minimum segment evidence score of 100 was used.

## Tumour mutational burden calculation

TMB was calculated by dividing the number of on-target, nonsynonymous somatic variants in each sample by the targeted panel territory in megabases (SureSelect v4 or v5). Variants were restricted to the panel regions with 100-bp interval padding. Nonsynonymous events were defined from SnpEff ANN consequence annotations to include nonsense, nonstop, splice-site, frameshift, in-frame insertion, and in frame deletion alterations^53^.

## Mutant-allele tumour heterogeneity calculation

Mutant-allele tumour heterogeneity (MATH) score was calculated as defined below, where is the median variant allele fraction for on-target, nonsynonymous somatic variants in that sample and MAD is the median absolute deviation (MAD) of VAFs, defined as

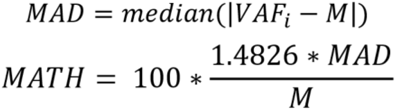

MATH scores were reported only when at least 10 qualifying variants were available^54^.

## Fraction of genome altered calculation

FGA was calculated by dividing the total nonredundant number of panel-covered base pairs overlapping ClinCNV AMP or DEL segments by the total number of base pairs in the covered panel territory, using panel intervals padded by 100 bp^55^.

## Aneuploidy score

Aneuploidy score was calculated as the total number of chromosome arms classified as gained or lost in each sample^56^. For each chromosome arm, total copy number was estimated as the overlap-length–weighted median of ClinCNV segments and compared with the sample-level ploidy. Arms with total copy number greater than ploidy + 0.5 were classified as gains, those below ploidy − 0.5 as losses, and all others as neutral. Only samples with successfully generated arm-level copy-number calls were included; samples with empty or unavailable ClinCNV CNA files were excluded.

## GISTIC analysis

GISTIC analysis was performed independently for relapse (n=104) and non-relapse (n=143) patient cohorts using input segment files derived from ClinCNV combined-run coverage segmentation outputs. For each sample, autosomal and sex-chromosome coverage bins were converted to GISTIC segment means as

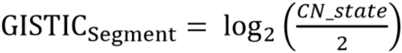

where *CN*_*state* is the per-bin ClinCNV coverage-derived copy-number estimate and 2 represents the diploid baseline. To reduce noise, bins exceeding the 90th percentile of within-sample variance were excluded. The remaining copy number profiles were smoothed using a centered rolling-median window of 71 bins, quantized in 0.30 log-ratio steps, and merged across genomic gaps of up to 300 kb. This preprocessing pipeline yielded a total of 28,254 smoothed segments for the relapse cohort and 37,069 smoothed segments for the non-relapse cohort.

To identify significantly amplified or deleted genomic regions, the smoothed segment data were processed using GISTIC2 (v2.0.23)^42^. The analysis was conducted utilizing the human genome assembly hg38 reference gene file (hg38.UCSC.add_miR.160920.refgene.mat).

For comparison with spatial transcriptomic *inSituType* data, this process was repeated with a few minor changes. The cohort was restricted for only the 25 total samples that were also analysed subsequently through spatial transcriptomics. copy-number profiles were smoothed across of 51 bins, quantized in 0.25 log-ratio increments, and merged across genomic gaps of up to 200 kb. Short segments spanning fewer than 20 bins were iteratively merged with the most similar adjacent segment. This preprocessing yielded 8,198 segments for the primary tumours and 2,672 segments for the metastatic tumours, for a total of 10,870 segments.

## Clonal inference with PyClone

Clonal inference was performed using PyClone-VI (v0.1.6) based on mutation-level input tables generated from the annotated somatic VCFs and copy-number states, which were assigned to each variant by overlaying ClinCNV segment call^5711^. Clones were defined by the PyClone-VI cluster assignments, and sample-specific clone abundance was quantified using the inferred cellular prevalence of the mutations within each cluster.

## Clone-dominance score

For each matched primary-metastatic tumour pair, PyClone-VI mutation-level cluster assignments and cellular prevalence estimates were used to quantify dominance among shared clones. Mutations detected in both samples were retained when their variant allele frequency was greater than 0.05 in both the primary and metastatic tumours and were grouped according to their primary-tumour PyClone cluster assignment. For each cluster *i*, *n_i_* is defined as the number of retained shared mutations and 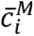 as their mean variant allele frequency. Only clusters containing at least two shared mutations (*n_i_* ≥ 2) were included. The weight of each cluster was calculated as

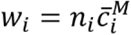

and the clone-dominance score (CDS) was defined as

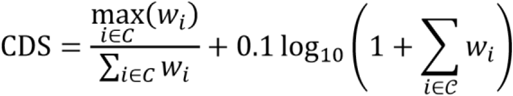

where *C*is is the number of retained shared clusters. The first term measures the proportion of the total cluster weight contributed by the dominant clone, whereas the logarithmic term modestly accounts for the total weighted shared mutational burden. Higher scores indicate stronger metastatic dominance by a single shared primary-derived clone. When no clusters met the inclusion criteria, the CDS was set to 0.

## Sequencing-based phylogenetic reconstruction of seeding trees

For monoclonal primary-metastasis pairs, evolutionary trees were reconstructed heuristically from PyClone-VI clusters. Primary-tumour clusters with more than one mutation were summarized by size, mean CCF, and the proportion of mutations shared with the metastasis. Clusters with at least 10 mutations were considered seed-eligible, and the cluster with the highest shared-mutation proportion was designated the seeding clone. Clusters were ordered by decreasing mean CCF. The seeding lineage was defined as the path from the trunk to the seeding clone. The metastatic root was attached to the most distal cluster on this lineage with at least 85% of mutations shared with the metastasis, or to the trunk if no cluster met this threshold. Metastatic subclones were organized using the same approach, based on mutations shared with the primary tumour.

## External-cohort acquisition and preprocessing (TCGA)

Copy-number alteration data for selected OncoCycle genes were analysed across cancer cohorts available through cBioPortal^36,58^. Only studies containing both gene-level copy-number and patient survival data were included. For OncoCycle deletion genes, analyses were restricted to homozygous deletions, and survival outcomes were compared according to alteration status within each cohort.

## Survival analysis

The primary endpoint was relapse-free survival, defined as the time from diagnosis to relapse or last documented follow-up and expressed in months. Patients without relapse were censored at the date of last follow-up. Survival analyses were restricted to primary tumours with complete genomic and clinical data. RFS distributions were estimated using the Kaplan–Meier method and compared across predefined genomic or clinical groups. Numbers at risk is defined as the number of patients who remained under follow-up and relapse-free immediately before each displayed time point.

## Cox regression and hazard testing

Time-to-event associations were modelled with Cox proportional hazards regression using lifelines (CoxPHFitter) with a penaliser of 0.01, fitting complete-case design matrices for each model. The main multivariable model included binary OncoCycle status, AJCC9 stage, tumour thickness group (<=2 mm, 24 mm, >4 mm), ulceration, TMB dichotomized at the cohort median, and FGA dichotomized at the cohort median. Hazard ratios (HRs) and 95% confidence intervals were taken directly from model coefficients. For multi-level covariates such as stage and thickness group, term-level significance was assessed by likelihood-ratio testing of the full model against a reduced model omitting that term.

## OncoCycle definition and scoring

OncoCycle was implemented as a binary six-gene panel comprising three functional biallelic loss genes (*CDKN2A*, *CDKN2B*, *TP53BP1*) and three copy-number gain genes (*CDK4*, *CD276*, *MCL1*). A gene was classified as having functional biallelic loss if it showed either homozygous deletion, or a loss-of-function oncogenic mutation together with heterozygous deletion or loss of heterozygosity. Gain-side events were defined from ClinCNV calls by the presence of amplification or gain affecting a target gene. For each patient, OncoCycle positivity was assigned if at least one loss-set gene met the functional biallelic loss criterion or at least one gain-set gene showed amplification/gain.

## Ternary plot computation

We analysed matched primary tumour and metastasis samples. For SNVs, OncoKB-annotated oncogenic variants were parsed and reduced to gene symbols. Ternary coordinates were calculated from the fractions of primary-private, metastasis-private and shared events across pairs. The 30 most recurrent oncogenic SNV genes are displayed in the final ternary plot. For CNVs, curated gene-level summaries from ClinCNV on target genes containing AMPs/Gains, and functional biallelic losses were derived and selected genes were plotted. A second CNV representation was generated by mapping broad hg38 cytobands and merging nearby bands on the same chromosome arm into broader regions.

## Identification of enriched CNA’s in relapsed

Cytoband annotations were collapsed to broad groups by removing sub-band designations (for example, 8q24.1–8q24.3 were merged as 8q24). Cytobands on the same chromosome arm, with the same direction of association and separated by no more than one band, were merged into larger regions. Each region was represented by the lead cytoband with the lowest Fisher’s exact P value. Regional false-discovery rates were calculated by applying the Benjamini–Hochberg correction to the set of lead cytoband P values. For volcano plots, regional significance was defined as a Benjamini–Hochberg false-discovery rate below 0.25^59^.

## Spatial Transcriptomics Analysis CosMx Assay

5μm FFPE tumour microarrays (TMA) were baked overnight at 60C and processed following the CosMx SMI Manual Slide Preparation for RNA Assays (MAN-10184-07). Briefly, target retrieval was performed at 100°C for 15 min and TMAs digested with proteinase K at 40°C for 30min. Probes (6125 plex) were hybridized to their targets in the tissues for 18hours. The TMAs were then stained with DAPI, CD45, PanCK, B2M and CD298 antibodies for cell segmentation staining. After washing, the flow cells were assembled and loaded onto the CosMx SMI instrument for imaging. Pre-bleaching profile configuration C (60 seconds) and Cell segmentation Profile A was chosen for the experiment with minor tweaks.

## Data Pre-processing – Quality Control (QC) and Filtering

Raw CosMX spatial transcriptomics data were exported from the AtoMX analysis platform as Seurat objects (one object per slide) and imported into R using the Seurat package (v5)^60,61^. Quality control was performed separately for each slide with identical filters. Retained cells had RNA counts between 50 and 10,000, a cell area between 40 and 600 µm², and were not flagged by AtoMx internal QC metrics. Cells localized at Field of View (FOV) borders were removed to mitigate technical artifacts arising from partial cell segmentation. Additionally, FOVs showing signal loss or failure at one or more barcode positions were excluded.

## Cell Typing

Dimensionality reduction was performed on the preprocessed data using Pearson-residual PCA (scPearsonPCA R package), which scales gene counts by their expected Poisson variance to produce a normalization-free embedding that is suitable for spatial transcriptomics data with variable capture efficiency^6222^. Data from all slides were merged without integration or batch correction prior to clustering, as the Pearson-residual normalization inherent to scPearsonPCA corrects for differences in sequencing depth. Variable genes were identified by applying the “FindVariableFeatures” Seurat function, selecting the top 2,500 variable genes to compute the PCA embedding. These embeddings were then used to construct a nearest neighbours graph. Cell clusters were identified using Seurat’s “FindClusters” function (Louvain algorithm) at a resolution of 1.2. For each cluster, marker genes were ranked by fold-change and fraction of expressing cells within versus outside the cluster. Cell type identities were then assigned based on cluster markers and closely related clusters were manually consolidated into broad cell type categories.

## Spatial Colocalization Analysis

The spatial co-occurrence between melanoma subclusters and immune cell populations was quantified using the Squidpy library in Python^63^. Slide-level coordinates were used as the spatial embedding. Cooccurrence scores were computed with “squidpy.gr.co_occurrence” across distance bins spanning 0– 300 µm in 50 µm increments. The co-occurrence score at a distance bin *d* between a reference cell type *X* and a query cell type *Y* is defined as the ratio of the observed number of type-*Y* cells at distance *d* from type-*X* cells to the number expected given the global proportion of type-*Y* cells in the dataset. A score greater than 1 indicates spatial enrichment, while a score less than 1 indicates spatial avoidance relative to random expectation.

## Differential Expression Analysis

Differential gene expression was assessed using the R package smiDE (spatial molecular imager Differential Expression), which is a framework developed for spatially-resolved transcriptomic data. For each gene, smiDE fits a negative binomial regression model^64^. The package accounts for segmentation bias by excluding genes whose expression in the target cell type is primarily explained by transcript overlap from neighbouring cells, and by including the aggregate expression of those neighbouring cells as a covariate in the model. To identify transcriptional changes from primary tumour to metastasis, tumour cells with paired primary and metastasis sections were analysed with tissue type (primary vs. metastasis) as the variable of interest. Differentially expressed genes were identified based on log2 fold change and FDR-adjusted p-values.

## Neighbourhood Enrichment Analysis

For every clone-labelled tumour cell, all other cells within a fixed radius of 50 µm were identified (RANN R package) and the composition of this neighbourhood was averaged across the cells per clone. Enrichment was assessed against a permutation null distribution generated by randomly shuffling the clone labels among the tumour cells (1,000 permutations). Cell positions and all non-tumour labels were held fixed, so that the clones are compared with one another. Per clone and cell type, a z score was calculated as the difference between observed and mean permuted neighbour fraction, expressed in units of the null standard deviation.

## Hypoxia Gradient Analysis

To quantify per-cell Hypoxia pathway activity, the UCell R package (v2.14.0) was used, which yields a score between 0 and 1 by ranking all genes within each cell and then deriving a Mann-Whitney U statistic over the ranks of the pathway genes^65^. The Hypoxia gene set was obtained from the MSigDB (release v2026.1.Hs) and restricted to the genes present on the CosMx 6k panel^6633^. The rank cutoff (maxRank) was set to the median number of genes detected per cell. For each cell, the distance to the nearest endothelial cell was determined by a k-nearest neighbour search with k=1 (FNN R package). Distances were grouped into eight predefined intervals, narrower near endothelial cells and wider in the more sparsely populated tail. The median Hypoxia UCell score was computed per bin.

## Gene Set Enrichment Analysis (GSEA) per clone

For each tumour subclone, all 6,175 CosMx panel genes were ranked by their average log2 fold change in that subclone versus all remaining subclones (Seurat’s “FoldChange” function). The ranked gene lists were tested against eleven gene sets (fgsea R package, v1.36.2)^67^, seven MsigDB hallmark sets and four melanoma differentiation subtype signatures^68^. The gene sets were restricted to genes present on the CosMx 6k panel. Enrichment scores were normalized by the mean enrichment score of random gene sets of the same size (normalized enrichment score, NES) to make gene sets comparable. P-values were corrected with the Benjamini-Hochberg procedure enrichment scores were normalized by the mean enrichment score of random gene sets of the same size (normalized enrichment score, NES), which renders the sets comparable with one another. These eleven sets were selected from a broader exploratory screen; p-values were corrected for multiple testing across all 55 subclone × gene set combinations shown (Benjamini-Hochberg) and are therefore conditional on this selection. Results are displayed as a dot plot in which colour encodes the normalized enrichment score (NES) on a scale symmetric about zero and dot size encodes −log10 of the adjusted p-value.

## Single Cell Copy Number Estimates

T cells and melanoma cells were extracted from all spatial transcriptomics samples (n = 25, 19 primary, and 6 met samples). A neighbourhood-based filtering step before SCNA, also CNV, inference. QuasiPoisson Pearson residual PCA was computed using scPearsonPCA. The resulting representation was used to construct a K-nearest-neighbour graph. Cells for which more than 20% of nearest neighbours (k = 50) were assigned to alternative cell type labels were removed. This filtering step removed 1.29% of melanoma cells and 1.53% of T cells.

CNV inference was performed separately for each sample using a procedure adapted from inSituCNV, a tool that infers copy-number variation from image-based spatial transcriptomics data^69^. Melanoma cells from each sample were used as test cells, while T cells pooled across all samples were used as the diploid reference population. Gene expression was summarized across genomic regions and compared with the Tcell reference to infer large-scale copy-number gains and losses in melanoma cells.

CNV inference was run independently for each sample. For cohort-level visualization, sample-level CNV results were concatenated to generate the final CNV heatmap. To reduce visual imbalance caused by samples with very large melanoma-cell numbers, samples containing more than 1,200 melanoma cells were randomly downsampled to 1,200 melanoma cells for heatmap visualization only. Samples with 1,200 or fewer melanoma cells were shown in full.

## Comparison with NGS Panel-Sequencing CNV calls

NGS CNV data from ClinCNV was imported and overlapped with the CNV bins generated by inSituCNV. inSituCNV CNV bins overlapping fewer than five NGS panel-sequencing probe regions were excluded from the validation analysis. Bins for chr6 also removed as T cells reference can create artificial loss in CNV at chr6. For each NGS panel-tested probe region, we first mapped the probe region to the inSituCNV CNV bin with the largest genomic overlap. inSituCNV CNV bins with less than 5 tested probe regions were excluded from analysis. Each tested region was then assigned a CNV change based on the ClinCNV CNV estimate at the midpoint of that tested probe region. The NGS panel CNV change was calculated as:

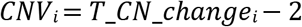

where *T*_*CN*_*c*ℎ*ange* is the absolute tumour copy-number estimate from ClinCNV. For each retained CNV bin, the panel-derived CNV change was calculated as the length-weighted average of all mapped tested regions:

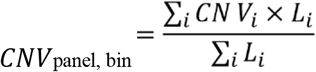

where *CNV_i_* represents the panel-sequencing CNV change for overlapping segment *i*, and *L_i_* represents the length of overlap between segment *i* and the CNV bin.

For each sample, CNV bins were assigned to one of three categories based on the panel-sequencing-derived CNV change: gain, neutral, or loss. Bins with panel-sequencing CNV change greater than 0.01 were classified as gains, bins with CNV change less than −0.01 were classified as losses, and bins with absolute CNV change less than or equal to 0.01 were classified as neutral. This threshold was used to remove extremely small nonzero CNV changes; 11 of 7,148 non-zero bins were assigned to the neutral category by this criterion.

For each sample and CNV-state category, transcriptomics-inferred CNV values were averaged across all bins assigned to that category. The corresponding panel-sequencing CNV values were averaged across the same bins. Thus, each sample contributed up to three data points, corresponding to gain, neutral, and loss categories; fewer data points were included when a sample lacked bins in one or more categories. Mean transcriptomics-inferred CNV values were plotted against mean panel-sequencing-derived CNV changes, and the association between the two measurements was assessed using Spearman correlation.

## CNV-defined clonal structure in patient FO13343

For patient FO13343, inSituCNV results for primary and metastatic samples were further analysed to characterize CNV-defined clonal structure using single cell based metrics instead of bulk sequencing. Cells were clustered based on their inferred CNV profiles using Leiden clustering on the inSituCNV results^70^.

CNV clusters were reviewed based on their genome-wide CNV profiles. In the primary sample, one small subclone containing 217 cells was merged with another clone because of similarity in CNV structure. In the metastatic sample, one small cluster containing 21 cells was removed from downstream clonal analysis because of small size and uncertainty on contamination.

Evolutionary trajectories were inferred by comparing CNV profiles across primary and metastatic subclones. Candidate clonal relationships were manually curated based on shared CNV events, clonespecific CNV gains or losses, and overall similarity of genome-wide CNV structure.

