## Supplementary figures and images for "Genomic correlates of metastatic competence and progression in human melanoma"

### Suppl. Figures

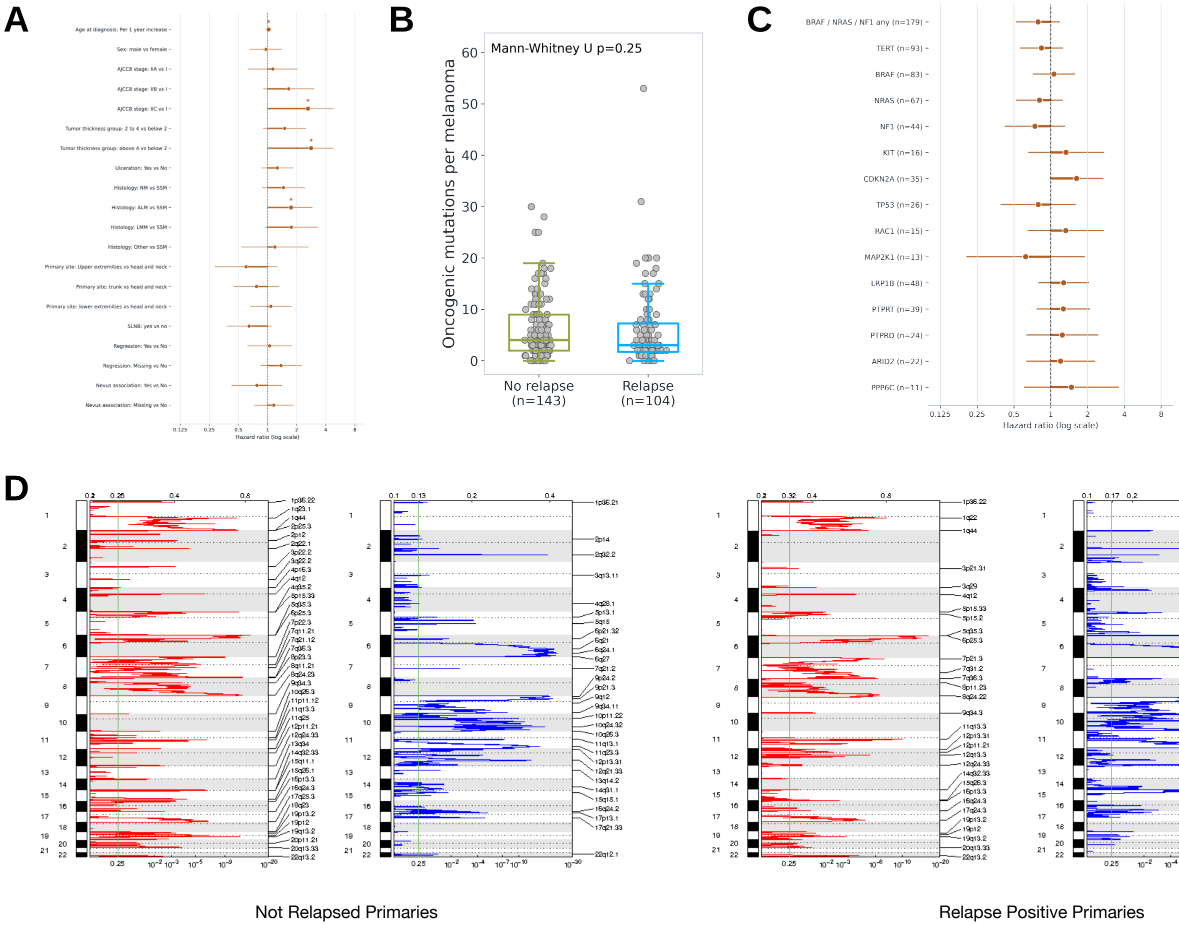


**Supplemental Figure 1**

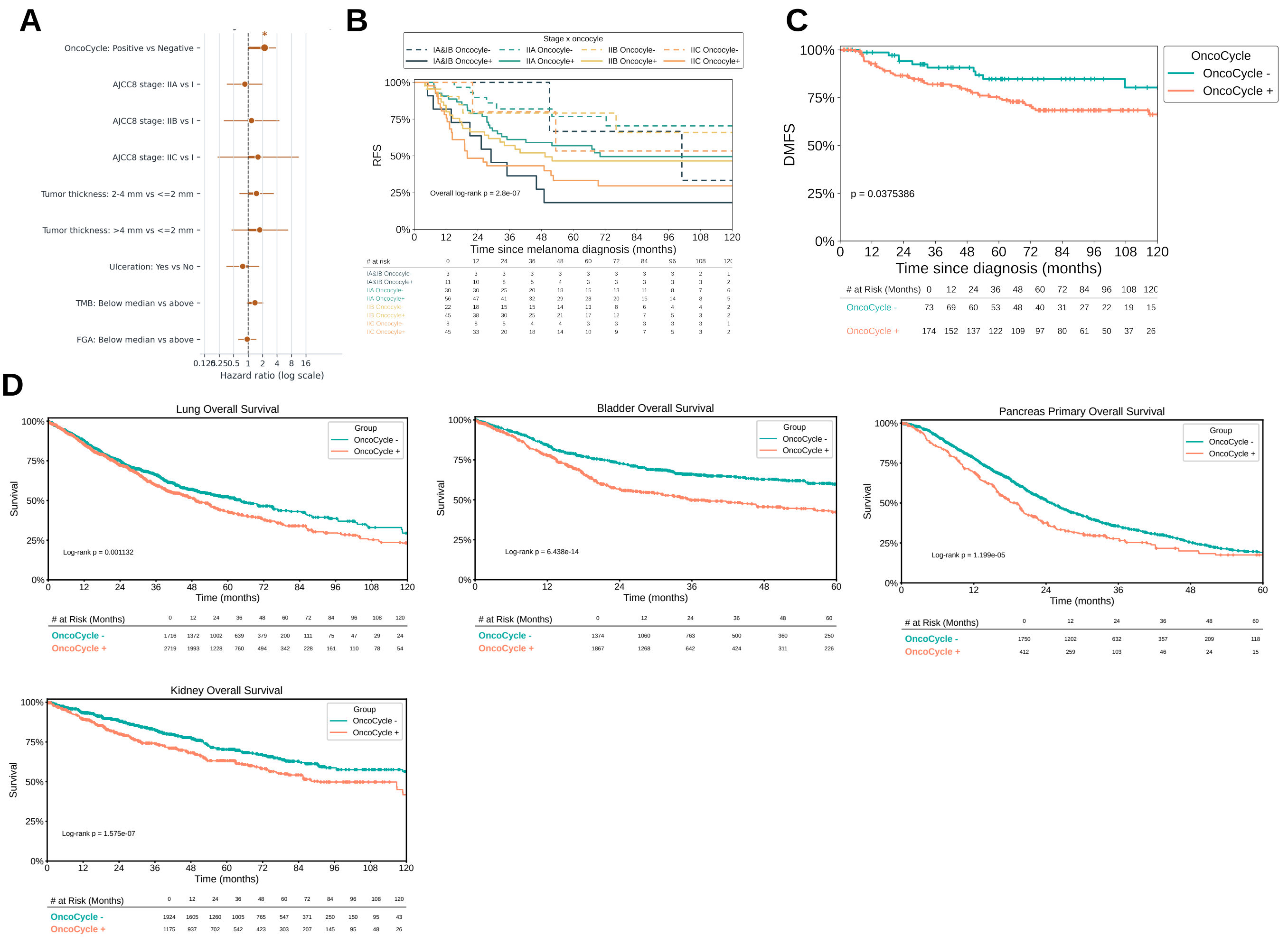


**Supplemental Figure 2**

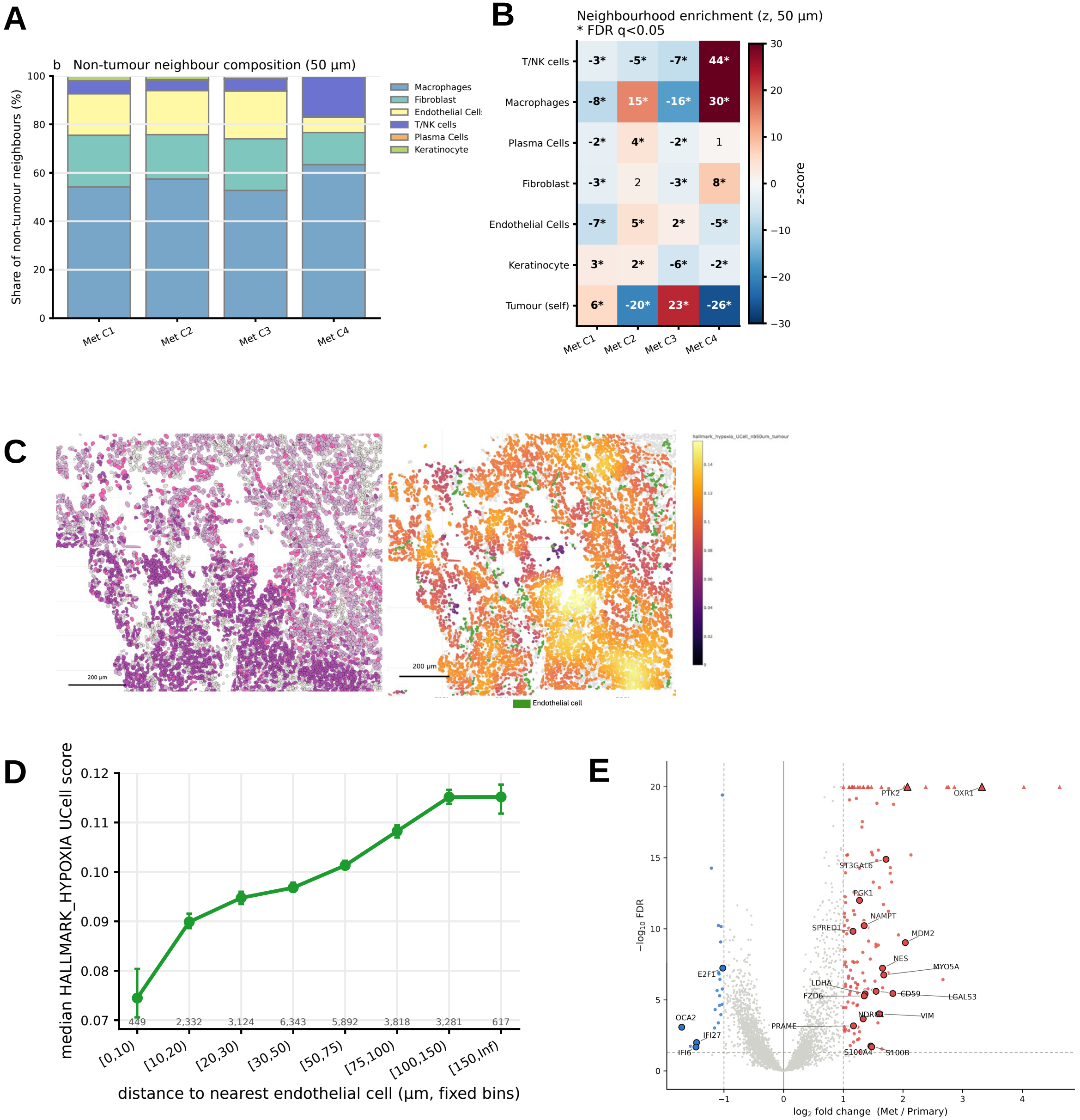


**Supplemental Figure 3**
